# FRS2-independent GRB2 interaction with FGFR2 is not required for embryonic development

**DOI:** 10.1101/2023.03.23.534012

**Authors:** James F Clark, Philippe Soriano

## Abstract

FGF activation is known to engage canonical signals, including ERK/MAPK and PI3K/AKT, through various effectors including FRS2 and GRB2. *Fgfr2^FCPG/FCPG^* mutants that abrogate canonical intracellular signaling exhibit a range of mild phenotypes but are viable in contrast to embryonic lethal *Fgfr2^-/-^* mutants. GRB2 has been reported to interact with FGFR2 through a non-traditional mechanism, by binding to the C-terminus of FGFR2 independently of FRS2 recruitment. To investigate if this interaction provides functionality beyond canonical signaling, we generated mutant mice harboring a C-terminal truncation (T). We found that *Fgfr2^T/T^* mice are viable and have no distinguishable phenotype, indicating that GRB2 binding to the C-terminal end of FGFR2 is not required for development or adult homeostasis. We further introduced the *T* mutation on the sensitized *FCPG* background but found that *Fgfr2^FCPGT/FCPGT^* mutants did not exhibit significantly more severe phenotypes. We therefore conclude that, while GRB2 can bind to FGFR2 independently of FRS2, this binding does not have a critical role in development or homeostasis.

## Introduction

Fibroblast Growth Factor (FGF) signaling plays an integral role in development, driving numerous cellular processes including proliferation, differentiation, and cellular adhesion (Clark and Soriano, 2022; Ornitz and Itoh, 2022; Ray et al., 2020; Ray and Soriano, 2023). FGFs are a family of secreted proteins that bind to and activate their cognate FGF receptors (FGFRs), which are receptor tyrosine kinases (RTKs). The mammalian FGF signaling family consists of fifteen canonical FGF ligands and four canonical FGFRs (Ornitz and Itoh, 2015; Ornitz and Itoh, 2022). Upon activation, FGFRs recruit multiple effectors to engage downstream intracellular signaling pathways, including ERK/MAPK and PI3K/AKT (Brewer et al., 2016).

Both *Fgfr1* and *Fgfr2* are necessary for early development (Ciruna and Rossant, 2001; Deng et al., 1994; Yamaguchi et al., 1994; Yu et al., 2003). Deletion of *Fgfr1* results in embryonic lethality at peri-implantation while deletion of *Fgfr2* results in lethality at midgestation on a 129S4 genetic background (Brewer et al., 2015; Kurowski et al., 2019; Molotkov et al., 2017). We have previously investigated how these FGFRs engage signaling to drive developmental processes by introducing point mutations that ablate the recruitment of specific effectors. Interestingly, the most severe combinatorial *Fgfr1* and *Fgfr2* alleles, *FCPG* (which eliminate binding of <u>F</u>rs2, <u>C</u>rk, Shb/<u>P</u>lcγ, <u>G</u>rb14), prevent the activation of all downstream canonical signals, but do not recapitulate the null alleles. Homozygous *Fgfr1^FCPG/FCPG^* embryos develop until at least E10.5, while *Fgfr2^FCPG/FCPG^* mice are viable. The large disparity between the FCPG and null alleles for both receptors indicates that partial functionality remains in both the *Fgfr1^FCPG^* and *Fgfr2^FCPG^* alleles (Brewer et al., 2015; Ray et al., 2020; Ray and Soriano, 2023).

Growth factor receptor-bound protein 2 (GRB2) is a cytosolic adaptor protein that plays a significant role in mediating downstream RTK activity. It consists of two SH3 domains and an SH2 domain. The SH2 domain of GRB2 specifically binds to phosphotyrosine residues on activated RTKs, while the SH3 domain binds to proline-rich regions on other signaling proteins such as Son of Sevenless (SOS). GRB2 recruits SOS to the plasma membrane, where it interacts with the small GTPase RAS, leading to the activation of downstream kinases in the MAPK pathway (Lowenstein et al., 1992; Rozakis-Adcock et al., 1993). Canonically, GRB2 is recruited to FGFRs via the adaptor protein FGF receptor substrate 2 (FRS2) (Kouhara et al., 1997).

It has also been shown that GRB2 can bind directly to the C-terminus of FGFR2, prior to ligand-dependent FGFR2 activation. This interaction regulates the phosphorylation state of FGFR2 by modulating the interaction of FGFR2 and the phosphatase SHP2, independently of the GRB2-dependent activation of ERK/MAPK via FRS2. Deletion of the ten terminal amino acids of FGFR2 was shown to abolish GRB2 recruitment to the intracellular domain of FGFR2, identifying the site of interaction (Ahmed et al., 2010; Ahmed et al., 2013; Lin et al., 2012). To determine if this novel function of GRB2 is involved in the residual activity of the *Fgfr2^FCPG^* allele *in vivo,* we generated mice harboring a deletion of the last ten amino acids of FGFR2 to prevent the recruitment of GRB2. On its own, the truncation (T) does not have a significant effect on development. When combined with a previous signaling allele *(Fgfr2^FCPG^),* we find that homozygous *Fgfr2^FCPGT/FCPGT^* mice display the same phenotypes as *Fgfr2^FCPG/FCPG^* mice, with little difference between the two. We conclude that the direct interaction between FGFR2 and GRB2 via the C-terminus does not have a required role in development.

## Results and Discussion

GRB2 has been reported to bind directly to FGFR2, interacting with phosphorylated Y812 and the last 10 amino acids of the C terminus. To determine if GRB2 binds to our signaling mutant allele, *Fgfr2^FCPG^*, we overexpressed *Fgfr2^FCPG-3xFlag^* in NIH3T3 cells. Using immunoprecipitation, we found that GRB2 was bound to FGFR2^FCPG-3xFlag^ even in the absence of FRS2 binding (Fig1A). Additionally, we found that GRB2 binds to wild-type FGFR2^WT-3xFlag^ in unstimulated conditions, as previously reported (Ahmed et al., 2010).

**Figure 1.**
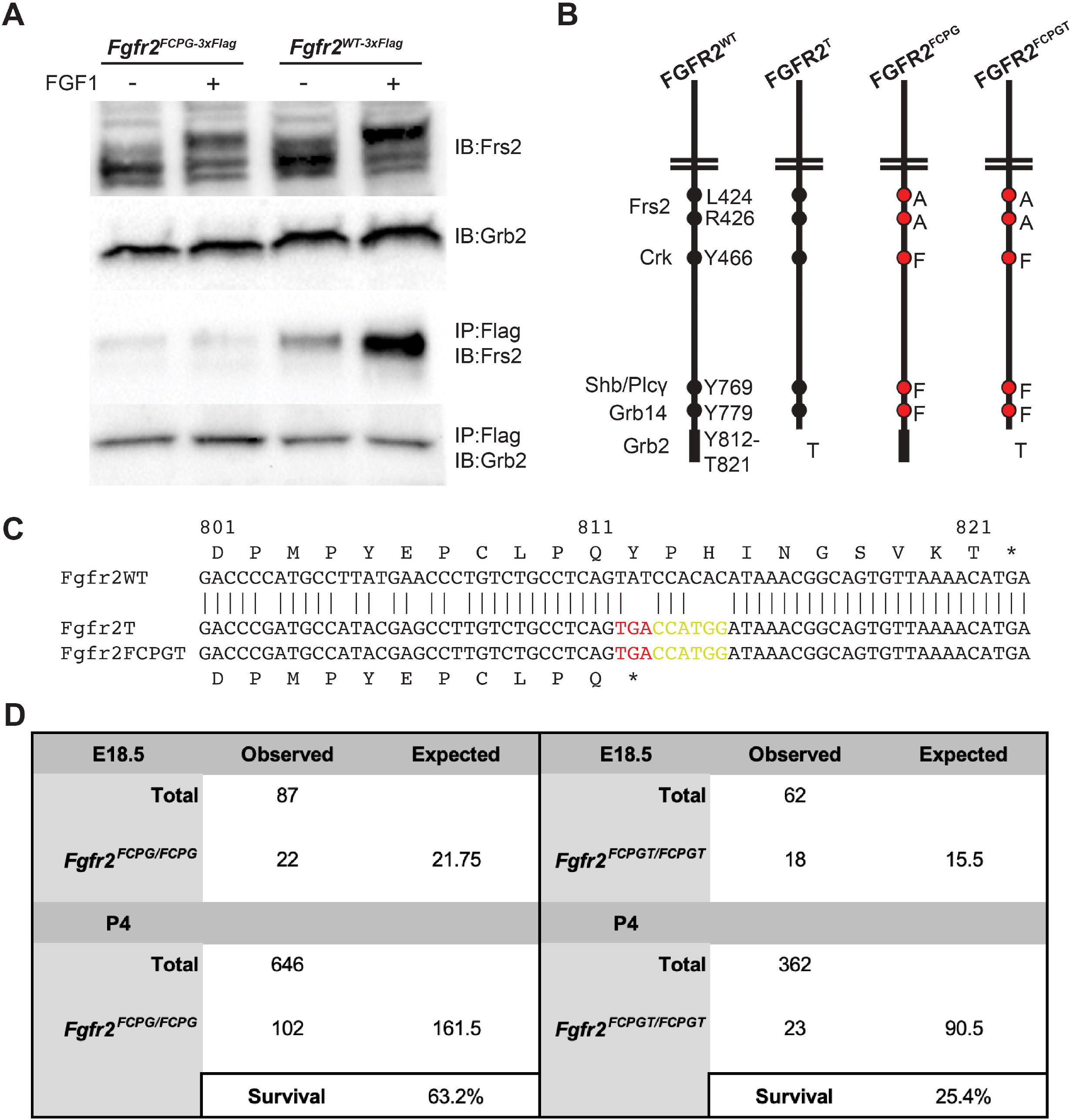
FCPG and FCPGT animals display similar phenotypes during adulthood and development. A) GRB2 is bound to FGFR2 in the absence of FRS2 in NIH3T3 cells expressing *Fgfr2^FCPG-3xFlag^*. GRB2 is also bound to wild-type *Fgfr2^WT-3xFlag^* during starvation conditions. The top two rows depict whole cell lysates, while the bottom two rows depict elution following immunoprecipitation with anti-Flag magnetic beads. Gel image is cropped to highlight changes. Full gel image is available in supplemental materials. B) Using 2C-HR-CRISPR, both *Fgfr2^T^* and *Fgfr2^FCPGT^* alleles were created to analyze the effects of GRB2 binding to the C-terminus of FGFR2. The most severe combinatorial allele, *Fgfr2^FCPGT^*, carries mutations to prevent the binding of FRS2, CRK, PLCγ, and GRB14, in addition to the C-terminus truncation. C) Sequencing of the C terminus of *Fgfr2* alleles. *Fgfr2^T^* and *Fgfr2^FCPGT^* both contain an early stop codon in place of Y812. An NcoI cut site (CCATGG) was also introduced after the stop codon to facilitate genotype differentiation. Raw sequencing data available in supplemental materials. D) Both *Fgfr2^FCPG/FCPG^* and *Fgfr2^FCPGT/FCPGT^* exhibit partial perinatal lethality; however, *Fgfr2^FCPGT/FCPGT^* does have a greater reduction in survival, p<0.001. Data for *Fgfr2^FCPG/FCPG^* is from (Ray et al., 2020).

We next examined if this interaction influences *Fgfr2* function during development. To impede GRB2 binding, we introduced an early stop codon at the *Fgfr2* locus via 2C-HR-CRISPR (Gu et al., 2018), truncating the 10 C-terminal amino acids (Fig1B, C). Heterozygous *Fgfr2^+/FCPG^* sperm was used to fertilize wild-type 129S4 oocytes; the fertilized zygotes were incubated until the two-cell stage after which they were injected with Cas9:sgRNA complexes and ssODN template and then transplanted into foster mothers. Of 23 offspring recovered, four were *Fgfr2^+/T^*, two were *Fgfr2^+/FCPG^*, and one was *Fgfr2^T/FCPGT^*, a 30% editing efficiency. Founders were then backcrossed to 129S4 animals to remove any potential off-target mutations.

On its own, the C-terminus truncation (T) displays no overt phenotypes. *Fgfr2^T/T^* animals are able to be maintained as homozygotes with no reductions in survival, litter size, or rearing capabilities, and adult mice appear indistinguishable from wild-type 129S4 animals (data not shown). We therefore decided to examine the T mutation in the sensitized context of our previously described *Fgfr2^FCPG^* allele (Ray et al., 2020).

A novel combinatorial allele, *Fgfr2^FCPGT^*, was obtained from the same 2C-HR-CRISPR experiment used to produce the *Fgfr2^T^* allele. The *Fgfr2^FCPGT^* allele harbors the new T mutation alongside the F, C, P, and G mutations that prevent the binding of FRS2, CRK, PLCγ, and GRB14, respectively (Fig1B, C). Two independent lines of mice, derived from two founders carrying the *Fgfr2^FCPGT^* allele, were initially analyzed to ensure consistency. Homozygous animals were recoverable and maintained from both *Fgfr2^FCPGT^* lines. Both *Fgfr2^FCPG/FCPG^* and *Fgfr2^FCPGT/FCPGT^* embryos were recovered in expected ratios at E18.5 (Fig1D), indicating that introduction of the C-terminal truncation in the sensitized *Fgfr2^FCPG^* background does not reveal the embryonic lethality seen in *Fgfr2* null embryos. As previously reported (Ray et al., 2020), *Fgfr2^FCPG/FCPG^* mice exhibit partial perinatal lethality, with roughly 60% surviving beyond P4. *Fgfr2^FCPGT/FCPGT^* mice also display partial lethality, but to a greater degree, with only about 36% surviving to adulthood, a significant reduction (p<0.001) compared to *Fgfr2^FCPG/FCPG^* animals (Fig1D).

During adulthood, both *Fgfr2^FCPG/FCPG^ and Fgfr2^FCPGT/FCPGT^* animals display similar phenotypes. Homozygous *Fgfr2^FCPGT/FCPGT^* mice are physically smaller than their wild-type or heterozygous littermates in both length (Fig2A) and weight (Fig2B). *Fgfr2FCPGT/FCPGT* mutants begin to exhibit periocular lesions around P15-P21 (Fig2C). This phenotype is also found in *Fgfr2^FCPG/FCPG^* mice and has been associated with defects in development of the lacrimal gland (Ray et al., 2020). Additionally, both *Fgfr2^FCPGT/FCPGT^* and *Fgfr2^FCPG/FCPG^* mice show partial penetrance of mild caudal vertebra defects, as evidenced by the presence of a kink in the tail (Fig2D).

**Figure 2.**
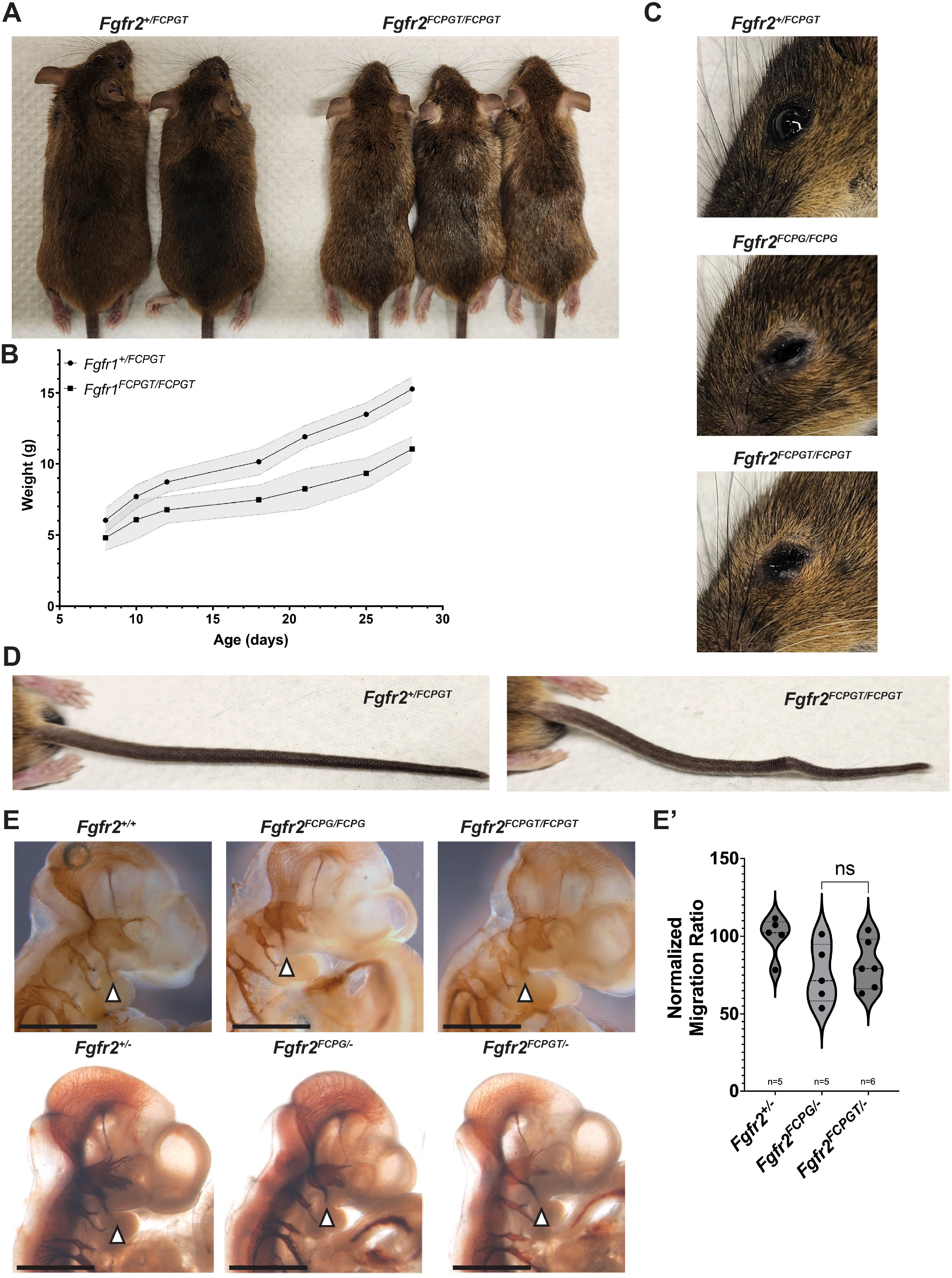
FCPG and FCPGT animals display similar phenotypes during adulthood and development. Homozygous *Fgfr2^FCPGT/FCPGT^* animals are viable but exhibit multiple phenotypes, including A) reduced size, B) weight, C) lacrimal gland impairment, and D) kinked tails compared to heterozygous littermates. E) Both *Fgfr2^FCPG/FCPG^* and *Fgfr2^FCPGT/FCPGT^* homozygous embryos displayed reduced migration of the trigeminal ganglion into the 1^st^ pharyngeal arch and is exacerbated in hemizygous embryos. However, there is no significant difference between the FCPG and FCPGT alleles. Images depict E10.5 embryos stained with anti-Neurofilament. E’) Comparison of migration ratios between hemizygous *Fgfr2^+/-^, Fgfr2^FCPG/-^*, and *Fgfr2^FCPGT/-^* E10.5 embryos. n.s., not significant. Line graph represents average weight and shaded area represents standard deviation. Violin plot represents median value on the dashed line and the dotted lines and tails represent quartiles.

Like *Fgfr2^FCPG/FCPG^* mutants, homozygous *Fgfr2^FCPGT/FCPGT^* mice can survive to adulthood and breed, albeit not as robustly as their heterozygous or wild-type littermates. As mentioned above, *Fgfr2^FCPGT/FCPGT^* mutants exhibit a reduced neonatal survival rate (Fig1D). Dead P0 pups that are recoverable lack a milk spot, indicating a failure to nurse. *Fgfr2^FCPG/FCPG^*, as well as hemizygous *Fgfr2^F/-^* and *Fgfr2^FCPG/-^*, were previously reported to have difficulty suckling, which might be due to cranial nerve defects (Ray et al., 2020). The trigeminal ganglion is a large nerve group that controls multiple motor functions in the face. During early development, the third branch of the trigeminal migrates into the first pharyngeal arch (PA) prior to E10.5, innervating the mandible, and is involved in suckling in neonates (Maynard et al., 2020). Reduced migration into the first PA was previously observed in *Fgfr2^FCPG/-^* embryos at E10.5 (Ray et al., 2020). We observed a similar phenotype in *Fgfr2^FCPGT^* mutants, as both *Fgfr2^FCPG/-^* and *Fgfr2^FCPGT/-^* embryos displayed reduced migration into the first PA (Fig2E), albeit with no significant differences between the two mutants (Fig2E’). A reduction in the ability to nurse is a probable cause for the partial neonatal lethality, as well as the reduced size of surviving mutants, as they would be outcompeted by their wild-type and heterozygous littermates.

There remains a large gulf between the *Fgfr2^-^* and *Fgfr2^FCPG^* alleles. We created the *Fgfr2^FCPGT^* allele to probe the residual function of the *Fgfr2*. On its own, the *Fgfr2^T^* allele appears to have no overt effects on embryonic or postnatal development, adult homeostasis, or reproduction. When combined with the previous signaling mutations, the *Fgfr2^FCPGT^* allele appears strikingly similar to the *Fgfr2^FCPG^* allele. The only significant difference observed between the two alleles was in neonatal survival. This finding suggests that the FGFR2 C-terminus, likely through GRB2 binding, contributes some activity to overall FGFR2 function, although this contribution must be minor as there was no difference seen in embryonic development. An alternative explanation is that the difference observed may be due to gradual genetic drift or changes in facility conditions over time, as the *Fgfr2^FCPGT^* data were collected in an independent, more recent cohort than the previously published *Fgfr2^FCPG^* data. As we did not see differences in any other phenotypes, the developmental impact of *Fgfr2* function seems to be negligible between the *Fgfr2^FCPG^* and *Fgfr2^FCPGT^* alleles. Questions remain as to how the *Fgfr2^FCPGT^* allele is able to maintain functionality in a manner sufficient for survival, while the *Fgfr2^-^* allele is lethal at midgestation. Further experimentation is needed to identify the unknown mechanisms by which *Fgfr2^FCPGT^* is compatible with life, without engaging canonical downstream signaling.

## Acknowledgements

We thank Leah Naraine for assistance with genotyping and cell culture, and Colin Dinsmore for discussions and comments on the manuscript. We thank Kevin Kelley and the Mouse Transgenic Core for their help in creating the novel *Fgfr2* alleles used in this work. This work was supported by F32 DE029387 (JFC) and R01 DE022778 (PS) from the National Institutes of Health (NIH)/National Institute of Dental and Craniofacial Research (NIDCR).

## Materials and Methods

### Animal husbandry

All animal experimentation was conducted according to protocols approved by the Institutional Animal Care and Use Committee of the Icahn School of Medicine at Mount Sinai (LA11-00243). Mice were kept in a dedicated animal vivarium with veterinarian support. They were housed on a 13 hr-11hr light-dark cycle and had access to food and water ad libitum.

### Mouse models and mutant generation

*Fgfr2^tm1.1Sor^*, referred to as *Fgfr2^-^*, and *Fgfr2^tm8.1Sor^*, referred to as *Fgfr2^FCPG^*, mice were previously described (Ray et al., 2020). *Fgfr2^T^* and *Fgfr2^FCPGT^* mice were generated by 2C-HR-CRISPR, as previously described (Gu et al., 2018). Briefly, heterozygous *Fgfr2^+/FCPG^* sperm was used to fertilize wild-type 129S4 oocytes, which were allowed to develop to the two-cell stage. Each blastomere was injected with preformed CAS9:sgRNA complexes and ssODN donor template. Edited embryos were subsequently injected into foster mothers and both *Fgfr2^T^* and *Fgfr2^FCPGT^* alleles were recovered. Heterozygous *Fgfr2^+/T^* and *Fgfr2^+/FCPGT^* animals were backcrossed to 129S4 at least six generations prior to analysis to remove any off-target editing. All mice were maintained on a 129S4 co-isogenic background. To differentiate the *Fgfr2^WT^* and the *Fgfr2^T^* or *Fgfr2^FCPGT^* alleles, oligonucleotides T_for_primer and T_rev_primer were used to amplify a 653bp fragment containing the mutation. PCR was followed by NcoI restriction digest to cleave the novel cut site induced alongside the T mutation (Fig1C).

### Oligonucleotides

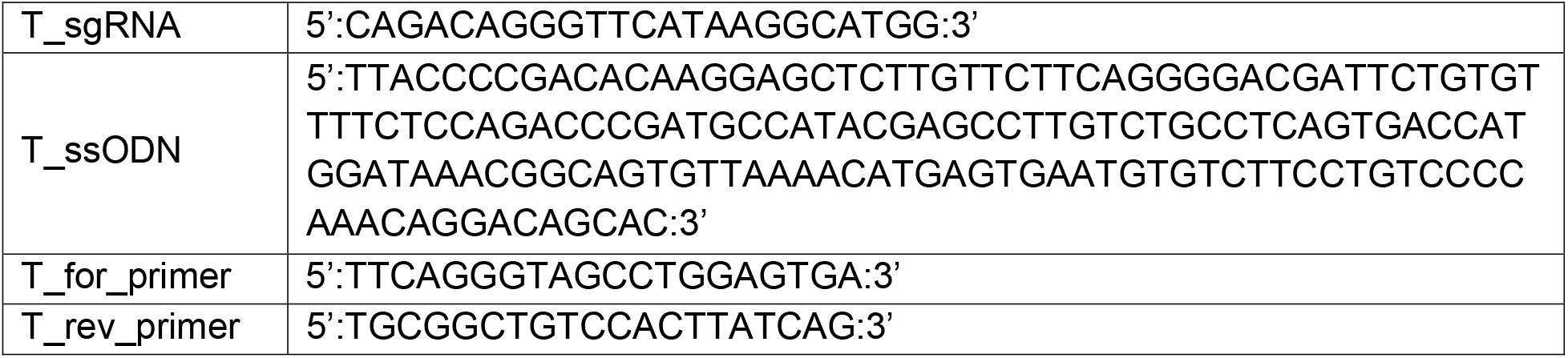

### Immunohistochemistry

E10.5 embryos were fixed in 4% PFA in PBS at 4°C, rinsed in PBS, then permeabilized in PBS with 0.5% Triton X-100 for 24hrs at 4°C. Neurofilament immunodetection was performed as previously described (Ray et al., 2020). Primary anti-neurofilament antibody (DHSB, 2H3) was used at a 1:20 dilution and anti-mouse HRP-conjugated secondary antibody (Jackson ImmunoResearch, 115-035-003) was used at 1:1000 dilution. Signal was developed using the ImmPACT DAB Substrate Kit (Vector Laboratories, SK-4105). Photographs were taken using a Nikon SMZ-U dissecting scope fitted with a Jenoptik ProgRes C5 camera.

### Coimmunoprecipitation

NIH3T3 fibroblasts were transfected with either pcDNA3.1-*Fgfr2c-Wt-3xFlag* or pcDNA3.1-*Fgfr2c-FCPG-3xFlag* via electroporation and stable lines were selected using the Neomycin resistance cassette co-expressed in the vector. Individual colonies were picked and cultured, and overexpression was verified by RT-qPCR and WB analysis. Cells were maintained in DMEM (Gibco, 11965118) supplemented with 10% HyClone FetalClone III (FCIII) serum (Cytivia, SH30109), 0.5X Penicillin/Streptomycin (Gibco, 15140122), 1X Glutamine (Gibco, 25030081), and 500ug/ml G418 (Gold Biotechnology, G-418). Prior to collection, cells were starved overnight for 18 hours in DMEM containing 0.1% FCIII, then treated with 50ng/ml FGF1 for 15 minutes. Cells were collected and lysed in NP-40/Digitonin lysis buffer containing protease and phosphatase inhibitors (Pierce, A32961) for 30 minutes at 4°C. 500 μg of lysate were incubated with anti-Flag M2 antibodies conjugated to magnetic beads (Millipore Sigma, M8823) overnight for 18 hours at 4C. Beads were collected and washed three times at 4°C followed by elution of bound protein using 4X Laemmli buffer heated to 95°C for 10 minutes. Binding of proteins was examined via Western Blot analysis using anti-Frs2 (SCBT, sc-8318) or anti-Grb2 (CST, 36344) used at 1:1000 dilution and anti-rabbit HRP-conjugated secondary (Jackson ImmunoResearch, 111-035-003) used at 1:10,000 dilution. Signal was developed using Immobilon Western (Millipore Sigma, WBKLS) and imaged using a ChemiDock MP (Biorad) imaging system.

### Statistical analysis

Statistical significance of neonatal survival was calculated using standard Chi-square analysis, with observed genotype frequencies compared to expected Mendelian frequencies. Chi-square analysis was also used to compare the observed genotype frequencies between the *Fgfr2^FCPG^* and *Fgfr2^FCPGT^* homozygous mutants.

The cranial nerve migration ratio was determined by dividing the length of the third branch of the trigeminal ganglion nerve by the length of the cranial nerve. Values were then normalized to the average ratio of *Fgfr2^+/-^* embryos. Statistical significance was calculated using One-way ANOVA with Bonferroni’s multiple comparisons test.

